# Unsupervised Neural Network-Based Image Stitching Method for Bladder Endoscopy

**DOI:** 10.1101/2024.09.24.614700

**Authors:** Zixing Ye, Chenyu Shao, Kelei Zhu

## Abstract

Bladder endoscopy enables the observation of intravesical lesion characteristics, making it an essential tool in urology. Image stitching techniques are commonly employed to expand the field of view of bladder endoscopy. Traditional image stitching methods rely on feature matching. In recent years, deep-learning techniques have garnered significant attention in the field of computer vision. However, the commonly employed supervised learning approaches often require a substantial amount of labeled data, which can be challenging to acquire, especially in the context of medical data. Both feature-based and unreliable supervised methods for cystoscopy image stitching are limited by their quality and the robustness of image stitching. This study proposes an unsupervised neural network-based image stitching method for bladder endoscopy that comprises two modules: an unsupervised alignment network and an unsupervised fusion network. In the unsupervised alignment network, we employed feature convolution, regression networks, and linear transformations to align images. In the unsupervised fusion network, we achieved image fusion from features to pixel by simultaneously eliminating artifacts and enhancing the resolution. Experiments demonstrated our method’s consistent stitching success rate of 98.11% and robust image stitching accuracy at various resolutions. Our method eliminates sutures and flocculent debris from cystoscopy images, presenting good image smoothness while preserving rich textural features. Moreover, our method could successfully stitch challenging scenes such as dim and blurry scenes. Our application of unsupervised deep learning methods in the field of cystoscopy image stitching was successfully validated, laying the foundation for real-time panoramic stitching of bladder endoscopic video images. This advancement provides opportunities for the future development of computer-vision-assisted diagnostic systems for bladder cavities.

## 1. Introduction

Bladder cancer is the ninth most common cancer worldwide, with approximately 430,000 new cases and 165,000 deaths annually [1]. Cystoscopy enables direct observation of intravesical cancerous features and serves as the gold standard for preoperative diagnosis and postoperative monitoring of bladder cancer [2]. However, cystoscopy is limited by its restricted field of view, making image stitching a key technique to address these issues

Computer vision (CV) and image stitching techniques aim to seamlessly combine multiple local images into a single complete image that is commonly used in scenarios such as panoramic generation. Conventional image stitching algorithms typically rely on feature point extraction, description, matching, and blending. In this regard, Behrens et al. [3] first proposed a process and method for panoramic fluorescence cystoscopy imaging. Subsequently, several researchers published solutions for mapping 3D scenes onto 2D planes in the human bladder for image stitching [4-6]. Building on previous studies, Bergen et al. [7-9] employed an approach based on prominent shared features. They registered and merged multiple consecutive frames from cystoscopy videos to create a larger field of view for bladder mapping, thus continually optimizing and ultimately developing software for processing real-time cystoscopy videos. However, their work was influenced by complex and variable noise within different bladder environments, weak texture, and blurring, leading to unstable stitching accuracy and significant degradation in stitching quality. To the best of our knowledge, bladder endoscopic image stitching is currently in its infancy, and only a few studies have been conducted. Thus, numerous challenges remain, including variations in illumination, weak textures, significant disparities, and poor stitching quality.

In recent years, with the rise of deep-learning technology, image stitching methods based on convolutional neural networks (CNNs) have become a research hotspot in CV. Deep learning models can automatically learn to extract features from raw images and achieve more accurate pixel alignment and blending during the stitching process, thereby significantly enhancing the quality and efficiency of image stitching [10]. Deep learning technology has been extensively explored in the field of bladder endoscopy for applications such as tumor identification [11], polyp classification [12], and hemorrhagic region semantic segmentation [13]. Deep learning can be categorized into supervised and unsupervised learning methods, based on the availability of labels [14]. However, to date, no unsupervised learning methods have been applied to bladder endoscopic image stitching, and supervised learning has been hindered by the challenge of obtaining stitching labels. A recent approach proposed the use of self-supervised learning to achieve image matching and alignment of bladder endoscopic images without relying on image labels [15]. However, the training complexity and alignment quality of this method remain suboptimal.

In this study, we propose a deep learning-based unsupervised stitching method for cystoscopy images, thereby aiming to enhance the robustness of stitching in various complex bladder scenarios and improve the quality of the stitched images. This method comprised an unsupervised alignment network and an unsupervised fusion network. In the former, we employed feature convolution and regression networks to estimate the homography matrix. Subsequently, we used tensor direct linear transformation (DLT) and a double-H transformer for image alignment. In the latter, we utilized pixelized deformation and refinement branches to learn image deformations and increase the resolution, thus achieving feature-to-pixel-level image fusion. By leveraging an available public cystoscopy image dataset, the strength of our approach is the proposal of an end-to-end stitching method for cystoscopy images that requires only two original images as input without the need for any labeled data during the training process.

## 2. Materials and Methods

### 2.1 Dataset

Obtaining bladder endoscopic images for training image stitching models is challenging, resulting in a scarcity of publicly available bladder endoscopic image datasets. Up to now, we have only found one publicly available bladder endoscopic image sequence [15] and a bladder phantom video created using 3D printing technology in literature [16] (used to simulate a real human bladder). In addition, six real human bladder videos were also collected in this paper to train the network. These videos were captured using different cystoscopes and encompassed various scenarios with arbitrary view angles: low overlap (<40%), medium overlap (40%–80%), and high overlap (>80%), with the difficulty of image stitching decreasing accordingly. They also featured different lighting conditions, blur levels, and textural strengths (dimmer, blurrier, and weaker textures further increase the stitching difficulty). The classification criteria for the overlap rate were developed based on references [17-19].

To create our training set, we split six real bladder videos and half of the bladder phantom videos into 2,248 frames distributed across different time intervals (5, 10, 15, and 20 frames). These frames were representative of diverse scenarios encountered during bladder endoscopy. They were used to generalize various real-world situations, as shown in Figure 1(a). The remaining videos were similarly divided into 521 frames to form the test set.

**Fig. 1.**
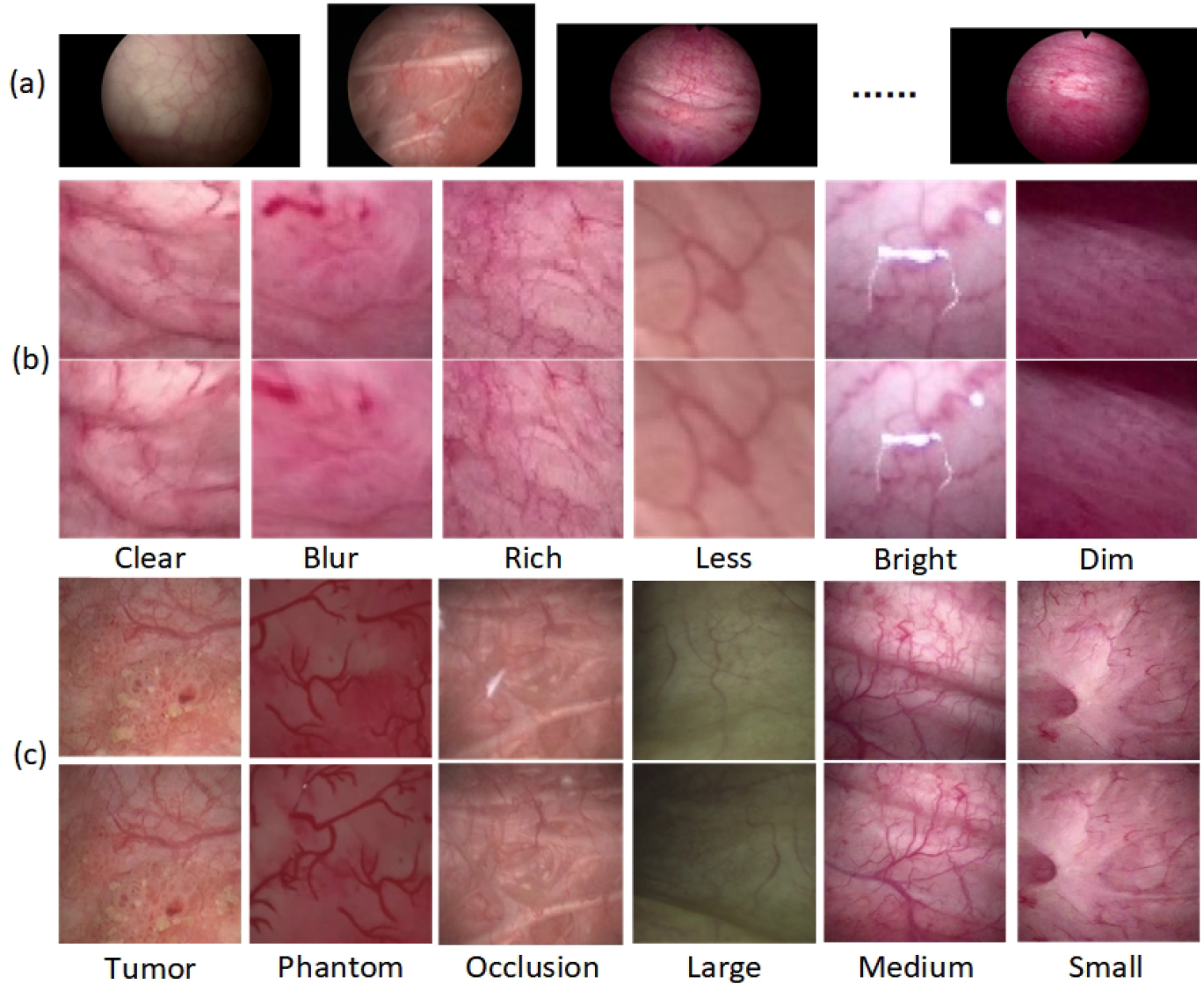
Dataset examples. (a) Different sequences of real bladder endoscopic images. (b) Augmented dataset with various features. (c) Original dataset with different scenarios

### 2.2 Preprocessing

Augmented dataset: The training of neural networks often requires large datasets. However, the available bladder endoscopic image sequences for training comprised only 2,248 images. To expand the dataset, we employed the data augmentation technique proposed by Bengio et al. [14]. Specifically, we randomly shuffled the existing real image sequences and selected 3/4 to create an augmented dataset consisting of 3,372 image pairs, as shown in Figure 1(b). Moreover, the augmented dataset was resized to 128×128 pixels.

Original dataset: After the initial training, we also needed a dataset for fine-tuning the pre-trained model. This dataset was sourced from the remaining 1/4 of the real image sequences. Subsequently, we applied proportional downsampling to each image and cropped the central region of interest, resulting in images resized to 512×512 pixels. From the 562 images, we manually selected image pairs with varying overlap ratios, which resulted in 477 image pairs. After random shuffling, these pairs constituted the original dataset for the unsupervised fine-tuning training, as illustrated in Figure 1(c).

### 2.3 Unsupervised Method

To achieve alignment and fusion of bladder endoscopic images, we introduced the stitching method proposed by Nie et al. [20], as depicted in Figure 2, which comprises two modules: an unsupervised alignment network and an unsupervised fusion network.

**Fig. 2.**
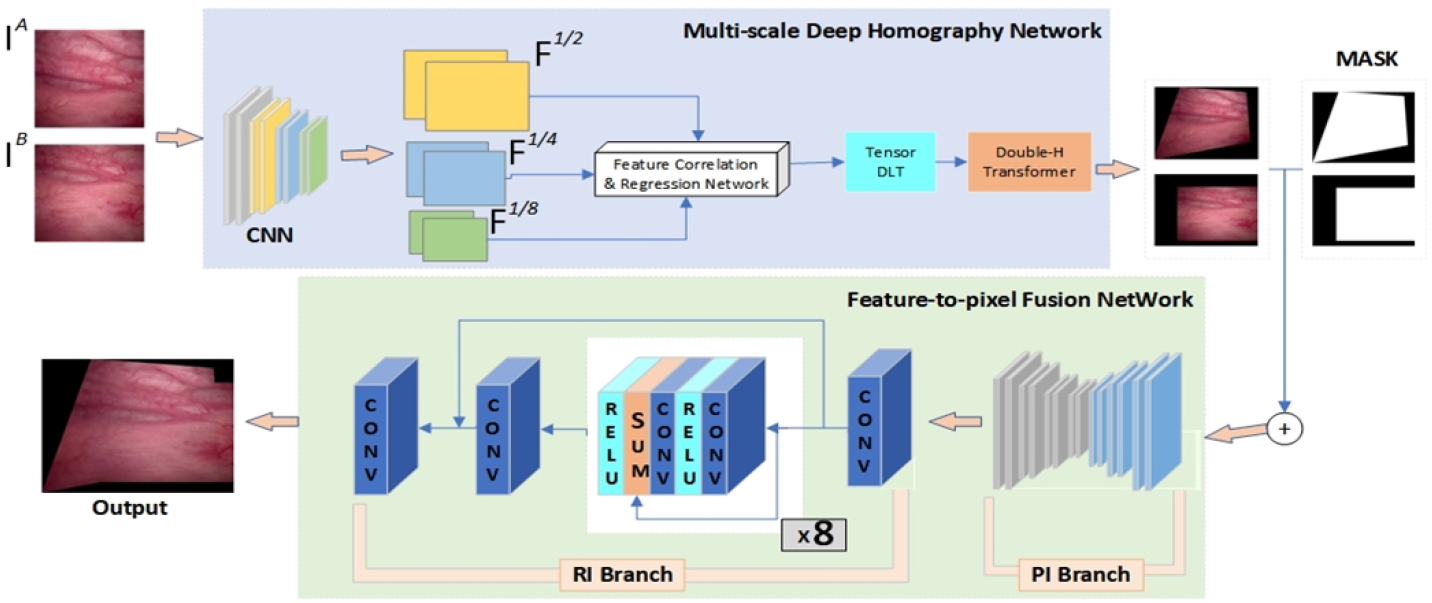
Unsupervised model architecture for cystoscopy images

In the first module, we employed a multiscale deep homography network to estimate the image homography when given a pair of bladder endoscopic images for stitching. Throughout this process, the network generated feature pyramids at 1/2, 1/4, and 1/8 resolutions of original image, which was employed in the subsequent analysis of feature correlation and regression. This enhanced the utilization of feature maps and expanded the receptive field of the bladder endoscopic images [14]. Subsequently, a tensor DLT was employed using the obtained homography matrix to warp the bladder images. These images underwent homographic transformations on a third black canvas, referred to as a double-H transformer, which enabled alignment between the two images. In this module, we adopted an ablation-based loss function to constrain the unsupervised homography network.

In the second module, the transformed bladder images from the first module were input into the unsupervised fusion network along with their respective masks. This network consisted of two branches: the pixelization deformation branch and refinement branch, which aimed to learn the deformation rules for image stitching and improve the resolution, respectively. The former involved downsampling the images and passing them through three pooling layers and three transposed convolutional layers; conversely, the latter included three individual convolutional layers and eight combination blocks, as shown in Figure 2. Finally, the outputs of the two branches were fused to generate a high-resolution pixel-wise stitched bladder image. This process enabled bladder image fusion from the feature level to the pixel level. In this module, we used content masks and seam masks to constrain the deformation rules of image fusion. The content mask ensured that the features of the fused image were close to the distorted image, while the seam mask ensured that the edges in the overlapping areas looked natural and continuous.

### 2.4 Training Setting

Two training phases were conducted for the unsupervised models as described herein. First, a deep homography network was trained on an augmented dataset for 120 epochs. Second, we fine-tuned the network parameters by an original bladder endoscopic dataset for 80 epochs. Throughout the training processes, we employed an unsupervised approach, utilizing only the reference and target images as inputs without the need for labels. An Adam optimizer with an initial learning rate of 0.0001, which decayed exponentially, was employed. The training and testing procedures were conducted using an NVIDIA GeForce RTX 3060 GPU.

### 2.5 Experimental Methodology

We conducted the evaluations in two stages: alignment and fusion. During the alignment stage, the robustness and quality of the alignment method were assessed. As representative of traditional feature-based methods, we selected Oriented FAST and Rotated BRIEF (ORB) [21], which is known for its speed, robustness, and effectiveness and is widely employed in conventional stitching approaches. Furthermore, we compared our method with local feature transformer (LoFTR) [22], a supervised deep learning-based alignment approach that represents current state-of-the-art supervised alignment methods. Our experiments were conducted at various image resolutions: 512×512, 256×256, and 128×128 pixels.

During the fusion stage, we conducted visual and quantitative analyses using the traditional Laplacian pyramid fusion [23] for comparison. The fused images were evaluated using metrics such as the peak signal-to-noise ratio (PSNR), root mean square error (RMSE), structural similarity index (SSIM), and total variation (TV). Through these metrics, our objective was to assess the visual quality and fusion performance of our bladder endoscopic image stitching method and compare it with traditional cutting-edge fusion approaches.

## 3. Results

### 3.1 Image Alignment

In this experiment, we categorized program execution errors that led to significant distortion, misalignment, and overlapping artifacts in the bladder images as “Errors,” as illustrated in Figure 3. These errors were considered failures. Our experiments involved a test dataset comprising 477 pairs of images, with 159 pairs in each of the large, medium, and small overlap categories. The statistical results of the alignment for each category are listed in Table 1.

**Fig. 3.**
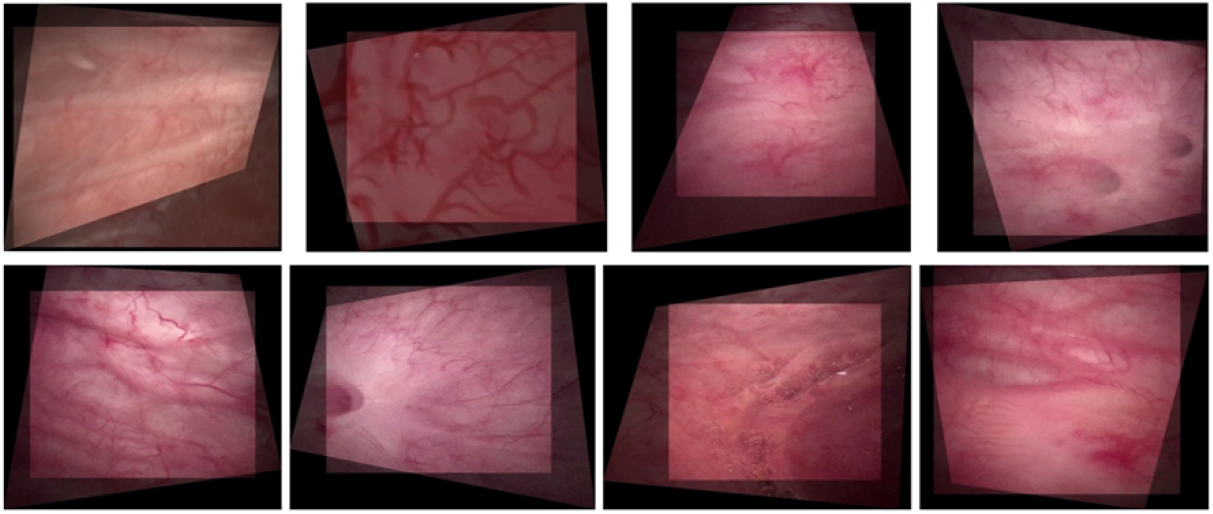
Images with distortion, misalignment, and overlapping artifacts

**Table 1:** Robustness testing of bladder image alignment, with a total of 477 test cases, including small overlap, medium overlap, and large overlap, each accounting for one-third of the cases.

| Input | Overlap grade | Metrics | ORB+RANSAC | LOFTR+RANSAC | Our |
| --- | --- | --- | --- | --- | --- |
| 512×512 | Large<br>(>80%) | Error | 6 | 0 | 0 |
|  |  | Failure | 46 | 2 | 3 |
|  | Middle<br>(40%–80%) | Error | 27 | 0 | 0 |
|  |  | Failure | 95 | 57 | 86 |
|  | Small<br>(<40%) | Error | 31 | 0 | 0 |
|  |  | Failure | 127 | 92 | 138 |
|  | - | Total | 332 | 151 | 227 |
|  | - | Success | 30.40% | 68.34% | 52.41% |
| 256×256 | Large<br>(>80%) | Error | 6 | 0 | 0 |
|  |  | Failure | 57 | 2 | 2 |
|  | Middle<br>(40%–80%) | Error | 30 | 0 | 0 |
|  |  | Failure | 104 | 59 | 86 |
|  | Small<br>(<40%) | Error | 45 | 0 | 0 |
|  |  | Failure | 109 | 93 | 139 |
|  | - | Total | 351 | 154 | 227 |
|  | - | Success | 26.42% | 67.71% | 52.41% |
| 128×128 | Large<br>(>80%) | Error | 19 | 0 | 0 |
|  |  | Failure | 70 | 2 | 3 |
|  | Middle<br>(40%–80%) | Error | 51 | 0 | 0 |
|  |  | Failure | 102 | 59 | 86 |
|  | Small<br>(<40%) | Error | 49 | 0 | 0 |
|  |  | Failure | 109 | 93 | 139 |
|  | - | Total | 400 | 154 | 228 |
|  | - | Success | 16.14% | 67.71% | 52.20% |

ORB: Oriented FAST and Rotated BRIEF [21], RANSAC: Random Sample Consensus, LOFTR: Local Feature Transformer [22]

We observed that the success rate of the feature-based ORB algorithm was only 30.04% at a resolution of 512 × 512 pixels and declined rapidly with increasing image blurriness, reaching only 16.14% at that of 128 × 128 pixels. However, both the supervised LOFTR method and our unsupervised learning method maintained a high success rates of approximately 67.71% and 52.41%, respectively, across different resolutions. Furthermore, as indicated by the blue highlights, our unsupervised method achieved an error count of only 2 or 3 in the category of large-overlap bladder endoscopic images at various resolutions, thereby maintaining a success rate of approximately 98.11%.

We visually assessed the quality using our bladder endoscopic image-alignment algorithm. As depicted in Figure 4, the feature-based ORB algorithm demonstrated significant distortion and severe image misalignment as the disparity increased and overlap decreased. Conversely, our unsupervised learning method, as illustrated in Figure 4(a) and 4(d), achieved visual results that can approach or even surpass those of LOFTR under various conditions, including small disparities and low illumination. In cases of moderate to large disparities, as shown in Figure 4(b-c), our method achieved alignment with only minor texture misalignment and small artifacts.

**Fig. 4.**
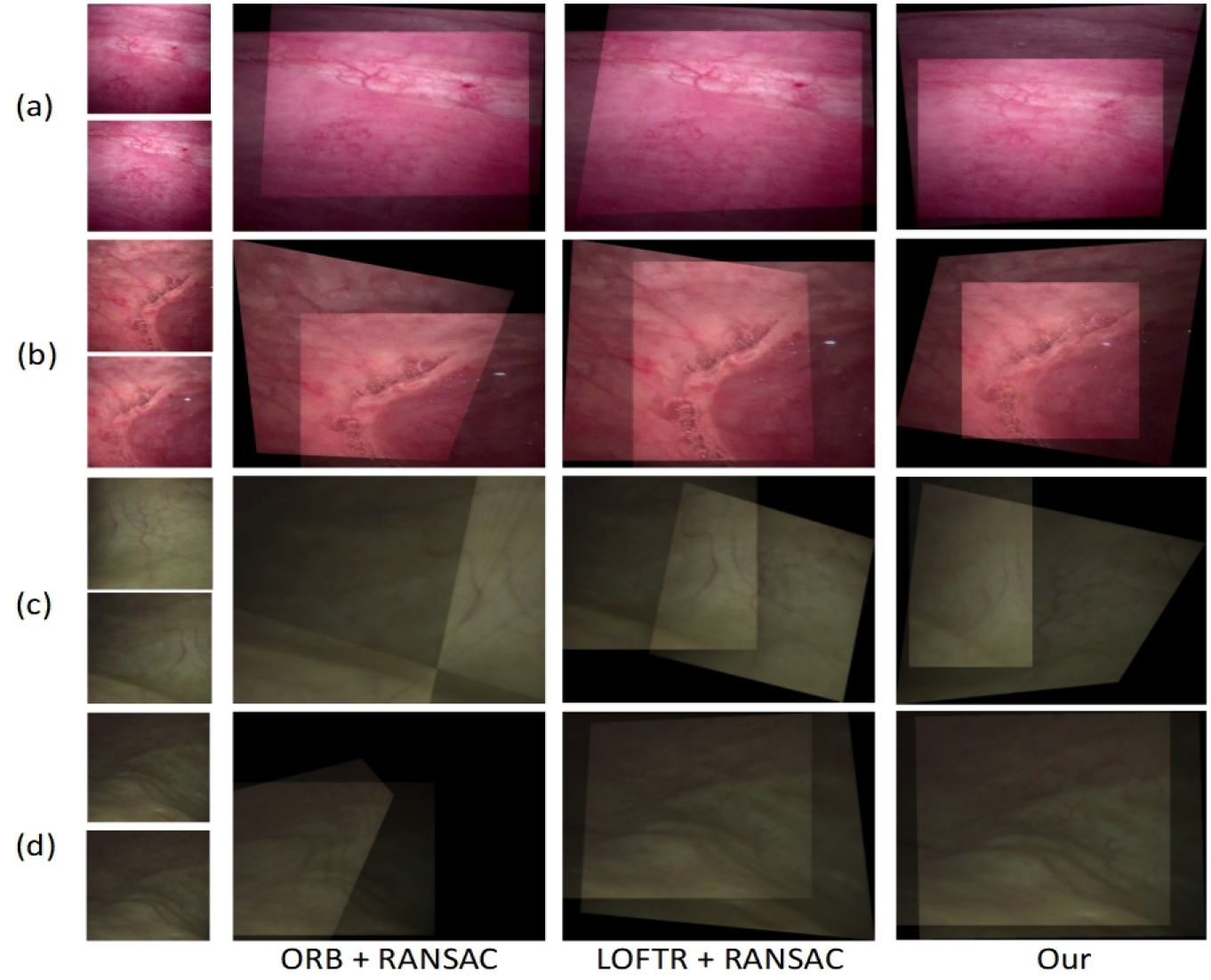
Comparison of alignment results. (a) Small disparity image (overlap rate > 80%). (b) Medium disparity image (overlap rate 40%–80%). (c) Large disparity image (overlap rate < 40%). (d) Image with extremely low visibility.

### 3.2 Image Fusion

In the second experiment, we conducted fusion tests on 477 image pairs from the test set and compared their visual quality with that of the traditional Laplacian pyramid method. As illustrated in Figure 5, our unsupervised approach effectively eliminated seam lines along certain boundaries. Furthermore, at the junction of the two images, our image fusion method ensured lighting consistency and reduced the visibility of artifacts and impurities in the bladder endoscopic images, as shown in Figure 5(a). Moreover, even after magnification, the fused images maintained a level of clarity similar to that of the original images.

**Fig. 5.**
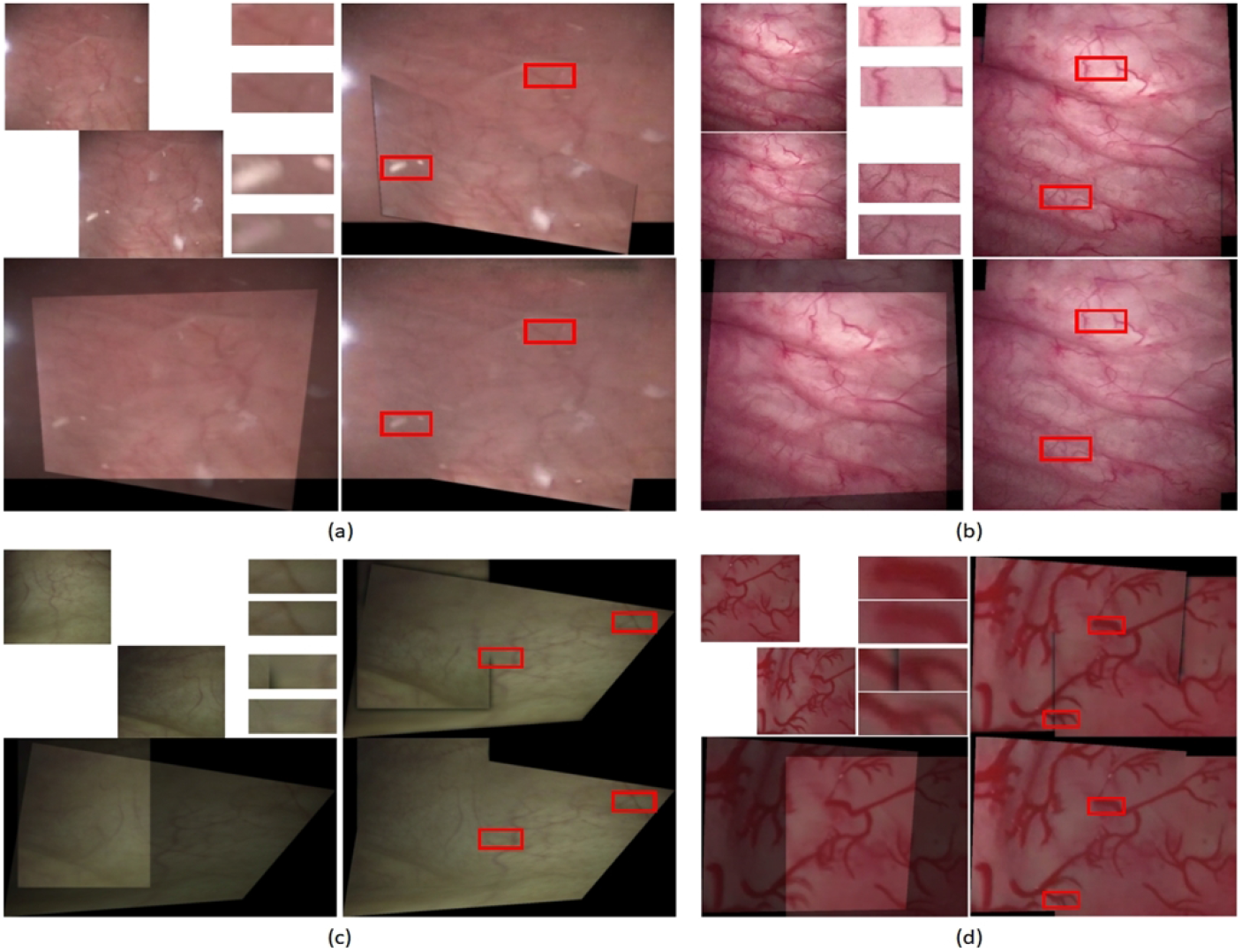
Comparison of fusion results. (a) Image with artifacts. (b) Image with clear and strong textures. (c) Image with dim and weak textures. (d) Bladder phantom image. In each scene, the top-left shows the input image pair, the bottom-left shows the aligned image, the top-right shows the fusion result using the traditional Laplacian pyramid method, the bottom-right shows the fusion result using our unsupervised method, and The middle image of the top-left and top-right corner shows the corresponding regions zoomed in by 2.5 times for comparison

Figure 6 presents the statistical performance analysis of the fused images generated by both methods compared with the original aligned images. It can be observed that the average range of PSNR falls within (15, 30), and the mean values of SSIM are close to 1 (Figure 6(a) and (c)). For most data sets, the PSNR curve of our method is slightly higher than that of traditional method, while the SSIM curve is slightly lower. However, it is evident that our fusion method consistently produces lower RMSE and TV values compared to the traditional method (Figure 6(b) and (d)). Particularly, the TV value is significantly lower than that produced by the Laplacian pyramid method (Figure 6(d)).

**Fig. 6.**
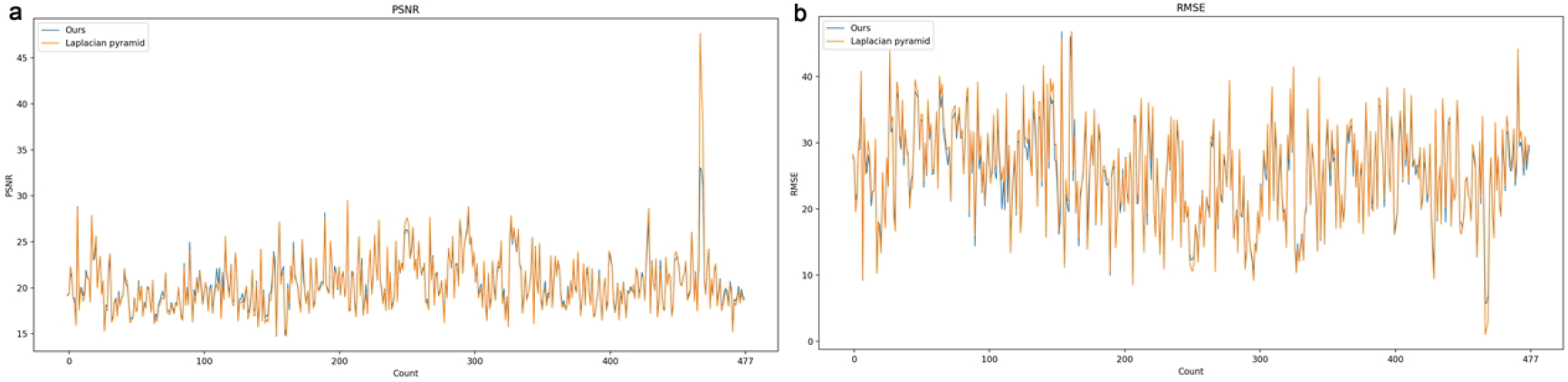
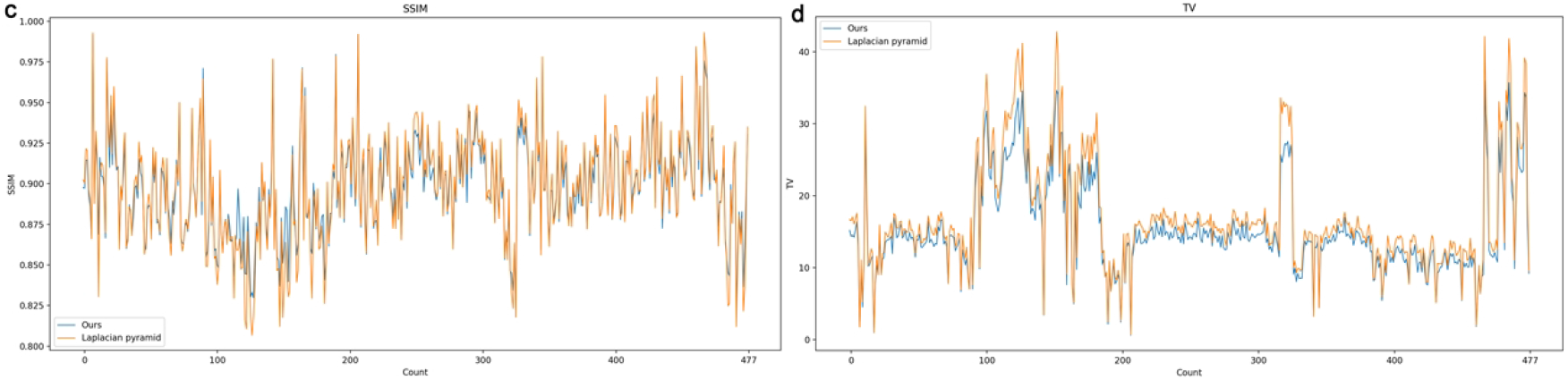
Performance line graphs of our unsupervised fusion and traditional Laplacian pyramid fusion on the test set. (a) Comparison of peak signal-to-noise ratio (PSNR) of the fusion images. (b) Comparison of root mean square error (RMSE) of the fusion images. (c) Comparison of structural similarity index (SSIM) of the fusion images. (d) Comparison of total variation (TV) of the fusion images

## 4. Discussion

The utilization of panoramic imaging for visualizing bladder lesions during cystoscopy has substantial clinical relevance, significantly enhancing the diagnostic efficacy and precision in detecting intravesical abnormalities. In this context, we proposed an unsupervised image stitching approach to facilitate the seamless amalgamation of bladder endoscopic images. Initial alignment experiments demonstrated the inherent susceptibility of the feature-based ORB algorithm concerning bladder endoscopic images. Notably, its performance rapidly deteriorated as image blurriness escalated, a trend consistent with the findings of Zhang et al. [21] in their investigations involving another endoscope. In stark contrast, both the supervised learning-based LOFTR method, and our unsupervised learning approach exhibited remarkable stability against decreasing resolutions. Although the performance of the supervised algorithm fell slightly below the 95% success rate attained in real-world scenarios [24], our unsupervised learning method closely approximated the outcomes of real-world testing [20]. Of particular significance is the comparable error rate of our unsupervised method in scenarios characterized by high overlap rates among bladder endoscopic images across diverse resolutions, which effectively matches the performance of the current optimal self-supervised algorithm in cystoscopy [15].

Based on the outcomes of this experiment in the alignment quality assessment, we inferred that the performance of the feature-based ORB solutions was sensitive to the quantity and distribution of feature points, thereby rendering their robustness inadequate across diverse scenarios. As the resolution decreases, our algorithm continues to exhibit steady alignment precision and resilience, effectively addressing the limitations inherent in conventional methodologies. In comparison with the supervised deep learning-based LOFTR method [22], our algorithm also demonstrates notable adaptability in tackling a range of complex scenes and disparities. Through a comprehensive exploration of visual perspectives and statistical success rates, our algorithm underscores that instances of model-based alignment failure occur predominantly in cases of moderate to large disparities. Importantly, our stitching effectiveness remained consistently high at smaller disparities, reaching a stable 98.11%. This achievement forms a solid foundation for the realization of panoramic stitching of continuous cystoscopy video frames, as initially proposed by Huo et al. [25].

In a subsequent fusion experiment, we offered a comparative analysis of the two techniques in terms of seamlessness, consistency, clarity, fidelity, noise, and artifacts, leveraging both intuitive visual observations and refined statistical metrics. As illustrated in Figure 5, the adaptability to varying intravesical image scenarios remains constrained, despite our diligent calibration of parameters, such as Gaussian kernel attributes, pyramid levels, and downsampling factors within the proposed Laplacian pyramid and its associated deformations [26]. This limitation is particularly notable when confronted with the complexities of texture and illumination aspects, which often result in discernible seamlines along specific boundaries. In contrast, our unsupervised approach effectively circumvented this concern. Although our methodology and the traditional approach [23] present comparable levels of visual quality, our impurity reduction and artifact mitigation capabilities are unattainable using the conventional method.

Furthermore, within the statistical assessment of the fusion experiment, while the comprehensive and average trends of the PSNR and SSIM depicted minor increments over those of the traditional method, discernible disparities were manifested in the RMSE and TV metrics. Notably, our fusion methodology consistently produced diminished values in these metrics relative to the conventional approach, underscoring the parity of our fusion model with the traditional approach in terms of image quality and structure. Specifically, our approach introduces a distinct advancement in image smoothness and texture preservation, which is congruent with the conclusions drawn from the subjective evaluations in the initial visual quality experiment. The resulting images exhibited enhanced smoothness, retained texture, and sustained luminance uniformity. The conspicuously reduced TV values further accentuate the relevance of this fusion model within the realm of endoscopic bladder imagery.

However, despite achieving a high success rate of approximately 98% in scenarios with small disparities in bladder endoscopy, our results are limited because of the scarcity of publicly available datasets specifically tailored for bladder endoscopy. Our dataset may consist of carefully curated samples. Thus, expanding the quantity and diversity of the datasets is necessary to improve the generalization ability of the model [27]. Furthermore, there is room for improvement compared with supervised learning-based methods, especially in cases involving moderate to large disparities and extremely low-texture regions. In addition, the lack of consensus and standardized evaluation metrics for image stitching and reconstruction in the field of endoscopy hinders the establishment of a universal benchmark for evaluating image stitching in bladder endoscopy [28].

## 5. Conclusion

To sum up, our method demonstrated robust image stitching accuracy at various resolutions and achieved an accuracy of 98.11% with a small disparity. Our method eliminates sutures and flocculent debris from cystoscopy images, presenting good image smoothness while preserving rich textural features, resulting in a more natural and detailed final result. Our application of unsupervised deep learning methods in the field of cystoscopy image stitching was validated, laying the foundation for real-time panoramic stitching of bladder endoscopic video images. This advancement provides opportunities for the future development of computer-vision-assisted diagnostic systems for bladder cavities.

## Statements and Declarations

### Author contributions

Z.Y.: Project development, Data collection, Manuscript writing, and Funding acquisition. C.S.: Methodology, Statistical analysis and Supervision. K.Z.: Methodology and References.All authors approved and contributed to the final manuscript.

### Funding

This work was supported by the Guangdong Basic and Applied Basic Research Foundation (2022A1515110036), National Natural Science Foundation of China (12302245), Natural Science Basic Research Program of Shaanxi Province (2023-JC-QN-0026), and Jiangsu Shuangchuang Talent Plan Project (JSSCBS20220943).

### Conflict of interest

The authors declare that they have no conflict of interest.

### Ethical approval

The study was approved by the Institutional Review Board and ethics committee and certify that the study was performed in accordance with the ethical standards as laid down in the 1964 Declaration of Helsinki and its later amendments or comparable ethical standards.

### Informed consent

was obtained from all individual participants included in the study.

## Notes

### Competing Interest Statement

The authors have declared no competing interest.

